# Finding Pathogenic nsSNP’s and their structural effect on COPS2 using Molecular Dynamic Approach

**DOI:** 10.1101/2020.10.12.333252

**Authors:** Ashish Malik, Kajal Pande, Abhishek Kumar, Alekhya Vemula, R Madhuri, Vivek Chandramohan

## Abstract

COP9 Signalosome Subunit 2 is a highly conserved multiprotein complex which is involved in the cellular process and developmental process. It is one of the essential components in the COP9 Signalosome Complex (CSN). It is also involved in neuronal differentiation interacting with NIF3L1. The gene involved in neuronal differentiation is negatively regulated due to the transcription co-repressor interaction of NIF3L1 with COPS2. In the present study, we have evaluated the outcome for 90 non-synonymous single nucleotide polymorphisms (nsSNP’s) in COPS2 gene through computational tools. After the analysis, 4 SNP’s (S120C, N144S, Y159H, R173C) were found to be deleterious. The native and mutated structures were prepared using discovery studio and docked to check the interactions with NIF3L1.On the basis of ZDOCK score the top 3 mutations (N144S, Y159H, R173C) were screened out. Further to analyze the effect of amino acid substitution on the molecular structure of protein Molecular Dynamics simulation was carried out. Analysis based on RMSD, RMSF, RG, H-bond showed a significant deviation in the graph, which demonstrated conformation change and instability compared to the wild structure. As it is known mutations in COPS2 gene can disrupt the normal activity of the CSN2 protein which may cause neuronal differentiation. Our results showed N144S, Y159H and R173C mutations are to be more pathogenic and may cause disease

## Introduction

COP9 Signalosome Subunit 2 (COPS2) is one of the important component of COP9 signalosome complex (CSN) that exists in eukaryotes. There are many synonyms that are in use for COPS2 like, TRIP15, SGN2 and CSN2. CSN is a conserved multi-protein complex (CSN1 to CSN8) that has seemingly evolved in parallel with the ubiquitin-proteasome system [1]. In a eukaryotic organism, CSN complex is associated to be as an essential protein complex that is involved in multiple cellular and developmental processes. Any deletion of CSN subunits can cause serious changes in the gene articulation profile and can cause early lethality of the multicellular organism [2–4]. The CSN complex was initially discovered as a eliminator of photomorphogenesis in *Arabidopsis thaliana [1]*. The eight mammalian complex subunits are classified according to the molecular weight S1 (57 kDa) to S8 (22 kDa) [5]. COP9 signalosome complex was considered as contaminating component in 26S proteasome preparation from erythrocytes by Seeger*et al.* and they later separated the two complexes [6]. The complex was cleansed biochemically from pig spleen by [5]. The CSN complex is a crucial regulator of ubiquitin (Ubl) conjugation pathway by mediating the deneddylation of the cullin subunits [7]. The subunits of CSN shows notable sequence homologies to elements of the lid subcomplex of the 26S proteasome which is formed by the assembly of 19S proteasome regulatory complex with the 20S catalytic core [8] and the translation initiation complex eIF3[6]. In eukaryotes, the eIF3 complex is involved in protein synthesis and ribosome biogenesis and it serves as a binding site for assembling other eIFsduring translation initiation [9]. Initially, Trip15/CSN2 was recognized as a thyroid hormone receptor (TR)-interacting protein and acts as a transcriptional corepressor [5]. In yeast two-hybrid screen, Trip15 is reported to interact with the thyroid hormone receptor and the retinoic acid X receptor in a ligand-dependent manner and the N-terminal region of subunit 2 (S2) is identical to Trip15[10]. Even though the mutual interaction between Trip15-nuclear receptor complex and CSN complex is obscure, indicating that the CSN complex assists in several signal transduction pathways [11]. COPS2 is a 51,597 Da protein with a sequence length of 443 amino acids. It also dissipates the interaction between OCT4 and CDK1 by segregating OCT4 and forming a COPS2/CDK1 complex thus hindering the inhibitory effect of OCT4 on CDK1 activation[12]. COPS2 (CSN2) shows transcriptional corepressor activity. The protein-protein interaction occurring inside the CSN complex (21-24) is stabilized by the proteasome COP9 complex initiation factor domain 3 in the C-terminal region of the Trip15/CSN whereas the N-terminal region of CSN2 has been reported to be adequate for the effector function of Trip15/CSN2. Trip15/CSN2 is associated with its binding partner through N terminal (<u>Akiyama et al. 2003</u>). The CS2 domain region (1-275) mediates interaction with NIF3L1, which is a highly conserved gene from bacteria to human. Although it is known that the mouse NIF3L1 gene encodes for a cytoplasmic protein (376 amino acids) and the expression of the protein was detected through mouse embryonic development, still the real-biological function of Nif3l1 is still unknown. In order to study the association of Trip15/CSN2 with Nif3l1, a wet lab study was carried out by *Akiyama and co* to exhibit that Nif3l1 protein could be translocated into nuclei by Trip15/CSN2 and it is involved in neural differentiation through down-regulation of Oct-¾ transcript which suppresses neurogenic genes. By the result of the study carried out by *Akiyama and co*, it seems likely that Nif3l1 and Trip15/CSN2 are associated and play an important role in neural differentiation [11]. NIF3L1 is engaged in retinoic acid primed neural differentiation of P19 embryonic carcinoma cells through cooperation with CSN2 [11]. In the complex COPS2 interacts with COPS1, COPS4, COPS5, COPS6, COPS7A or COPS7B directly[8, 13]. Notably, it also interacts with the ligand binding region of thyroid receptor, CUL1 and CUL2 [8]. Single nucleotide polymorphism is known as the most common type of variants occurring in human genome due to single nucleotide substitution [14, 15]. Non-synonymous SNPs can cause 50% of the disease that is related to inheritance [16]. SNPs can also disrupt the normal function of the protein and denature the structure by changing the stability, folding pattern and ligand binding site [16]. The amino acid changes at key sites within a protein may result in series of conformation changes, including the disintegration of salt bridges, alteration of interaction network, disruption in hydrogen bonds, which in turn may disturb the energy landscape [17]. Protein structure modeling methods have been extensively used for predicting the effects of disease-causing mutation on protein stability and protein-protein interaction [18].

The aim of the present study was to identify the most deleterious nsSNPs within COPS2 gene and their effect on protein structure. Protein-protein docking was done in order to determine the effect of amino acid substitution on the structure of COPS2 which will affect the interaction with Nif3l1. To check the interaction of COPS2 with NIF3L1, protein-protein docking was done. The mutations which are identified to have a positive interrelationship with pathogenicity and have high ZDOCK score were further investigated using MD simulations to determine the changes on its time-dependent physiological affinities and biochemical pathway alterations (<u>Sinha et al. 2018</u>). We utilized RMSD, RMSF, H-Bond, Gyration, in order to examine the motion trajectory and atomic interactions of native and mutant protein structures. Alteration in the protein structure can lead to a change in the binding site of protein which may affect the interacting proteins of COPS2 that may cause disease.

## Materials and Methods

### SNPs Data Collection

The data of human COPS2 gene were obtained from the NCBI database of dbSNP (www.ncbi.nlm.nih.gov/snp), with SNPs ID, amino acid positions, mRNA accession numbers, and protein accession number. The protein sequence of COPS2 gene was obtained from the UniProt database (UniProt ID: P61201) for further computational analysis. The workflow, for different tools analysis and databases, to identify the potential pathogenic SNPs of COPS2 gene are shown in Figure 1.

**Figure 1.**
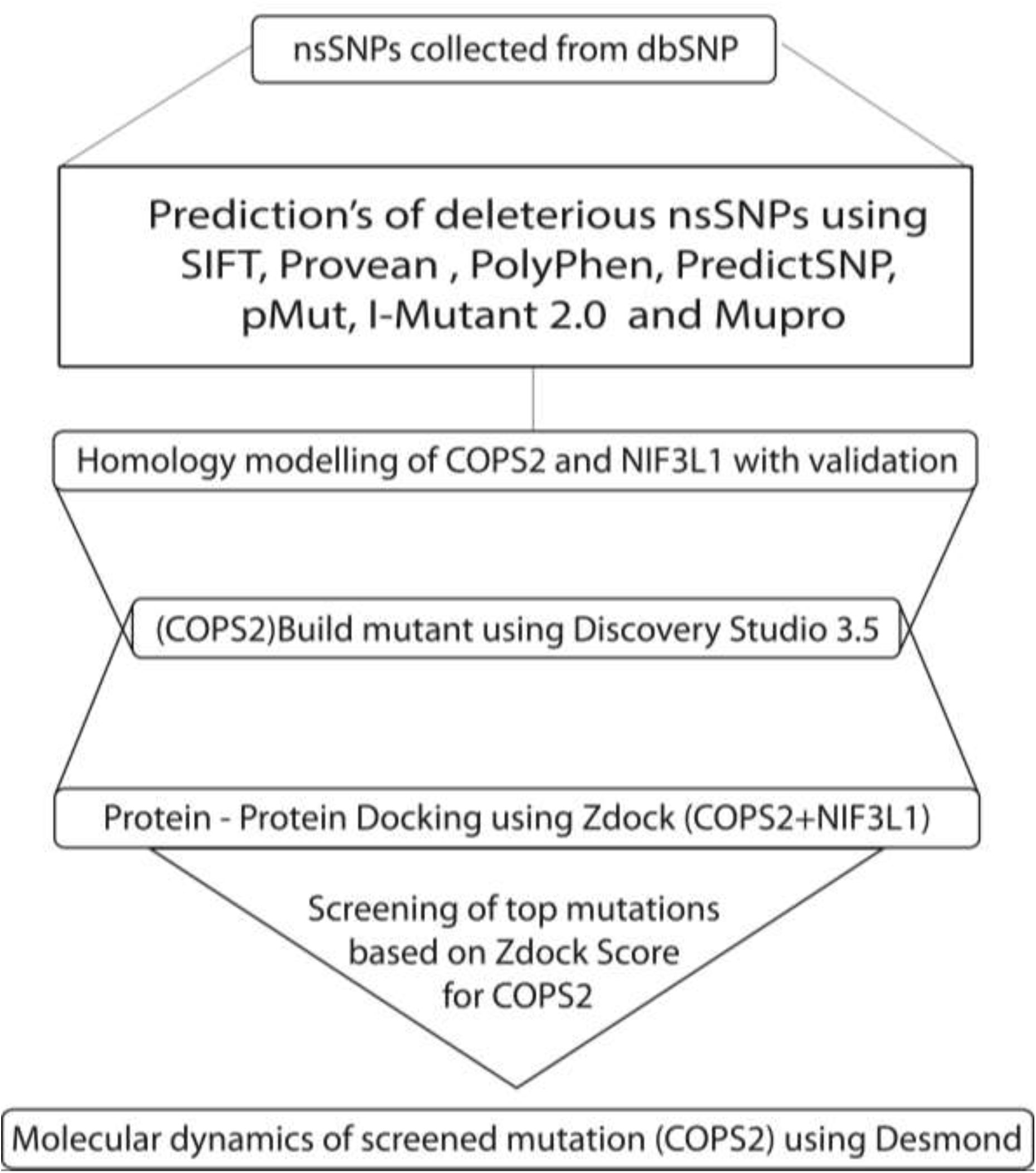
The flowchart depicting the overall process of identification and characterization of pathogenic SNPs in COPS2 along with structural and functional consequence analysis upon mutation.

### Evaluation of missense SNPs functionally

The function-based prediction of missense SNPs was predicted using 5 different tools, SIFT [18], PROVEAN [19], PolyPhen2 [20], PredictSNP [21], and PMut [21]. In these 5 tools, the query protein sequence is submitted in FASTA format with amino acid variants, i.e., mutational position and substitutions. While the stability-based prediction of missense SNPs was predicted using I-Mutant 2.0 [22] and MUpro [23]. In these tools, the query protein sequence is submitted in FASTA format with mutation position and substitute amino acid. The main reason behind the selection of functional tools and stability based tools to bottleneck the pathogenic mutations from the screened mutation database.

### Protein Modelling and Preparation of mutant structures

The crystal structure of COPS2 protein was obtained from PDB (Protein Data Bank), PDB ID = 4D10 (B Chain) [24]. The crystal structure has missing residue from position 181 to 191, henceforth SWISS-MODEL [25] was used to fill the missing residue using a template (PDB ID = 4WSN; B Chain) [26] with QMEAN significant Score of - 3.90 which is below the threshold of −4.0 and having a sequence identity of 100%. The COPS2 interacting protein NIF3L1 sequence (UniProt ID: Q9GZT8) was also modeled using SWISS-MODEL [25] using template (PDB ID = 2GX8, A chain) [27] as the best template with a sequence identity of 25.15% and with QMEAN score of −3.38, since there was no crystal structure present in the Protein Data Bank (PDB). After homology modeling, the modeled structures were validated using PROCHECK [28] for structural validation, stereo chemical properties, and Ramachandran plot analysis. Then the modeled structures of COPS2 and NIF3L1 were prepared at pH 5.4 and 7.5 in Discovery Studio 3.5 (Accelrys, Inc., San Diego, CA, USA.) as the parent structure was also crystallized at the same pH [27, 28]. Six mutated structures (R100W, S120C, N144S, N152K, Y159H, R173C) of COPS2 were prepared using Discovery Studio 3.5 with build mutant protocol (Accelrys, Inc., San Diego, CA, USA) and the Discrete Optimized Protein Energy (DOPE) were calculated [29, 30]. The QMEAN scores were also calculated for the mutated structures using SWISS-MODEL QMEAN tool [31] with the score of −3.29, −3.24, −3.28, −3.2, −3.27 and - 3.38 respectively. Which is all below the threshold of −4.0. therefore considered as a significant [32].

### Protein-Protein docking of COPS2 with NIF3L1

Protein-Protein interaction is highly significant because most of the biological activity depends on the specific recognition of proteins. The interactions between COPS2 with NIF3L1 structures were docked using Discovery Studio 3.5 ZDOCK protocol (Accelrys, Inc.). Where COPS2 was ligand protein and NIF3L1 was receptor protein, with angular step size of 6, Distance Cutoff of 10A^°^, RMSD cutoff of 10^A°^, Interface Cutoff of 10A^°^ and Maximum Number of Clusters was 100. The ZDOCK is a rigid-body protein-protein docking which uses Fast Fourier Transform Correlation technique and Critical Assessment of Prediction of Interactions (CAPRI) algorithms to delve the rotational and transitional space of a protein-protein system to create protein clusters in which top poses were screened using ZDOCK score [33].

### Molecular Dynamics

Schrödinger-Desmond (Desmond Molecular Dynamics System, D. E. Shaw Research, New York, NY) was used for the molecular dynamics simulation with Desmond v3.6 package with SPC water model. Orthorhombic periodic boundary conditions (PBC) was set up to specify the shape and size of the protein unit with buffering of 10A^°^. For neutralization of the system electrically, Na+/Cl− ions were added, pH was set to 5.4 and it was placed randomly in the solvated system. Molecular Dynamics Simulation was carried out in the NPT (Normal, Pressure, Temperature) ensemble using OPLS-AA (Optimized liquid Simulations –all atoms) 2005 force field parameters. The temperature and pressure kept at 300K and 1atm. respectively. The operation was followed by running in 150ns NPT production simulation in triplicate to confirm that protein was not stuck in local minima. After the molecular dynamics simulation, the structural changes in the monomer COPS2 protein molecule and mutant’s proteins were analyzed through RMSD, RMSF, SSE, Radius of gyration (Rg) and Hydrogen bonding [34–36].

## Results & Discussion

### SNP Screening

For COPS2 gene a total no. of 7440 mutations in *Homo sapiens* were found from preliminary search in dbSNP database where 90 SNPs were missense variant accounting 1.2 % of total mutations, these 90 SNP’s were subjected to Function based prediction tools (SIFT, PROVEAN, Polyphen2, PredictSNP, and PMut) and the following screened deleterious SNPs were then analyzed with Stability based prediction tools (I-Mutant & MUpro) for determining the structural stability of protein. (Supplementary Table 1)

### Identification of deleterious nsSNPs using function-based tools

Results of Function-based prediction tools (SIFT, PROVEAN, Polyphen2, PredictSNP, PROVEAN, PMut) and Stability based prediction tools (I Mutant 2.0, MuPro) are displayed in (Supplementary Table 2). SIFT classified 13 nsSNPs to be highly deleterious with a score ranging from (0.00–0.05) which are predicted to bring change in protein function. PROVEAN predicted 29 nsSNPs to be deleterious with the score ranging from (−2.55 to −6.54) with a threshold value of equal to or below −2.5 predicted to be deleterious with binary classification (i.e. deleterious vs neutral). PolyPhen 2.0 classified 14 nsSNPs to be ‘‘Possibly damaging’’ with scores ranging from (0.494-0.944) shows an impact on the protein function. PredictSNP shows 14 nsSNPs to be deleterious from other neutral residue showing the integrated result. PMut screened 18 nsSNPs to be in Disease condition with scores ranging from (0.51 to 0.82) predicting the alteration in protein functioning. In Mutant 2.0, 72 nsSNPs were predicted to decrease the stability of the protein and the remaining 18 nsSNPs were predicted to increase the stability of the protein based on the predicted ∆G score. MuPro server predicted 88 nsSNPs to be altering the protein structure due to decrease in stability. Among 90 nsSNPs in COPS2 gene functional and stability based tool predicted 12 nsSNPs in which 4 nsSNPs (S120C, N144S, Y159H, R173C) falls in the active domain (30 to 275 aa) which is interacting with NIF3L1 gene (Akiyama et al. 2003) the screened nsSNPs are tabulated in Table 1.

**Table 1:** Most pathogenic nsSNPs identified by function and stability based tools.

| Rs ID | Variant | SIFT | Provean | PolyPhen | PredictSNP | pMut | I-Mutant 2.0 ($\Delta G$) Kcal/mol | Mupro($\Delta G$) Kcal/mol |
| --- | --- | --- | --- | --- | --- | --- | --- | --- |
| <u>rs1361040987</u> | S120C | (Damaging)0.02 | Deleterious (-4.405) | Damaging (1.00) | Deleterious (0.755) | Disease (0.64) | Decrease (-1.66) | Decrease (-0.8601) |
| rs1480090014 | N144S | (Damaging) 0.25 | Deleterious (-4.975) | Damaging (1.00) | Neutral (0.602) | Disease (0.66) | Decrease (-0.990) | Decrease (-0.9907) |
| rs1317524775 | Y159H | (Damaging) 0.03 | Deleterious (-3.992) | Damaging (0.985) | Deleterious (0.605) | Neutral (0.49) | Decrease (-1.105) | Decrease (-1.1051) |
| <u>rs769174765</u> | R173C | (Damaging) 0.01 | Deleterious (-5.988) | Damaging (0.999) | Deleterious (0.869) | Disease (0.66) | Decrease (-0.19) | Decrease (-0.5031) |

### Protein modeling

The active domain region of COPS2 [30-275 aa] and NIF3L1 protein were modeled using SWISS-MODEL since the domain region [30-275] of COPS2 shows interaction with NIF3L1 [11]. Figure 2 Model [30 - 275 aa] (Green color) and Template [4WSN] (Red color) superimposed structure.

**Figure 2:**
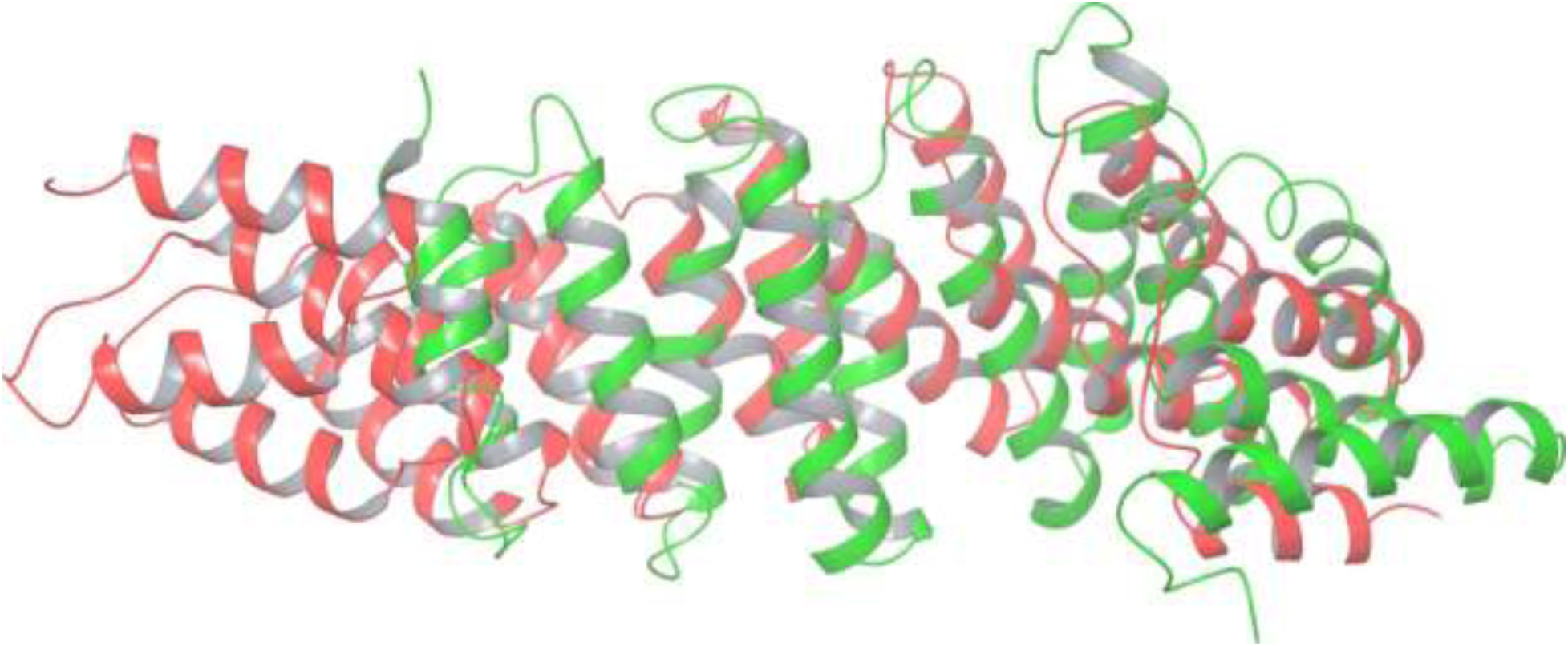
Superimposed Homology Modeled Structures (Red Color: Template and Green as a modeled structure.)

Structural validation of COPS2 and NIF3L1 was carried out using PROCHECK with Ramachandran plot. From Ramachandran plot Figure 3(A) it was found that COPS2 active domain region shows scoring pattern to be 198 (87.2%) residues in favored region with 24 (10.6%) residues to be in allowed region, 1(0.6%) to be in generously allowed region and 4 (1.8%) residues in disallowed region. For the structural quality of Homology modeled structure of NIF3L1 was resulted by Ramachandran plot Figure 3(B). Scoring pattern showing 274 (86.7%) residues are in favored regions, 38 (12.0%) residues are in allowed regions, 2 (0.6%) residues in generously allowed region and 2 (0.6%) in disallowed region.

**Figure 3.**
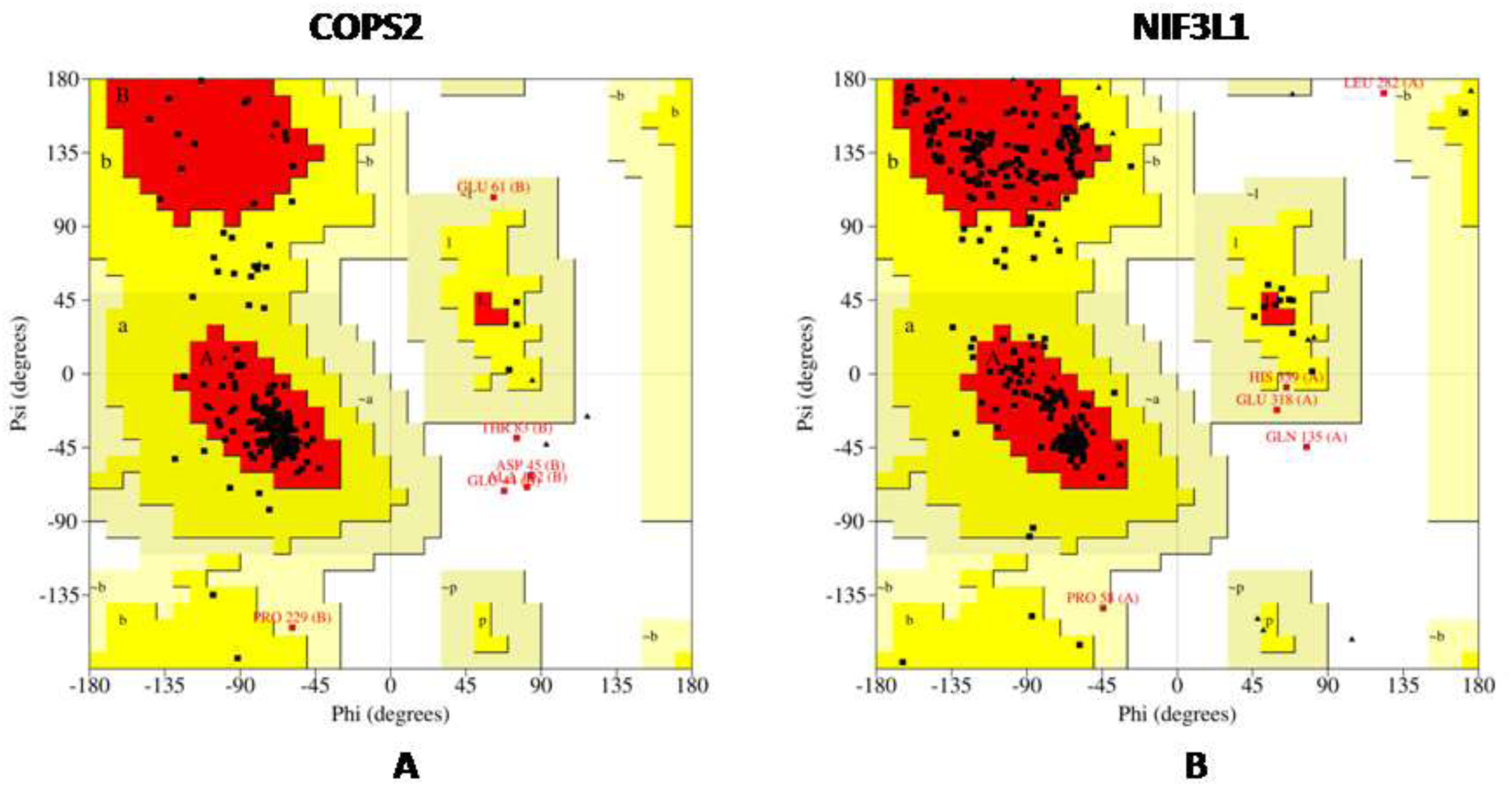
Quality and Structure assessment using Ramachandran plot for the modeled proteins. (A) COPS2 (B) NIF3L1.

Validation of above structures, COPS2 (Template) [4WSN] was superimposed with the Model [30-275 aa] Fig 2. Mutant structures were prepared using Discovery Studio 3.5 with build mutant protocol (Accelrys, Inc., San Diego, CA, USA) resulted, where Total Energy, Physical Energy, and DOPE score were calculated. Table 2

**Table 2:** Discrete Optimized Protein Energy (DOPE) values and mutation stability values of the mutated protein models.

| Mutations | DOPE Score |
| --- | --- |
| N144S | -26758.44141 |
| Y159H | -26520.50195 |
| R173C | -26797.23438 |
| S120C | -26863.17578 |

### Protein-Protein Docking Results

Focusing on protein-protein docking of the active domain of COPS2 with NIF3L1 homology modeled structure was taken [Template PDB ID:2GX8] (*Bacillus cereus*). NIF3L1 and domain region [30-275aa] of COPS2 (WT) were docked using Discovery Studio 3.5 ZDOCK protocol along with mutated structures (NIF3L1-S120C complex, NIF3L1-N144S complex, NIF3L1-Y159H complex and NIF3L1-R173C complex), where 2000 conformations resulted from which top conformation was selected on the basis of ZDOCK score. Only mutations (above 25>) were parameterized, 3 nsSNPs (N144S, Y159H, R173C) with ZDOCK score of 25, 25.14 and 25.66 respectively were selected for further analysis. The Protein-Protein docking was tabulated in Table 3.

**Table 3.**
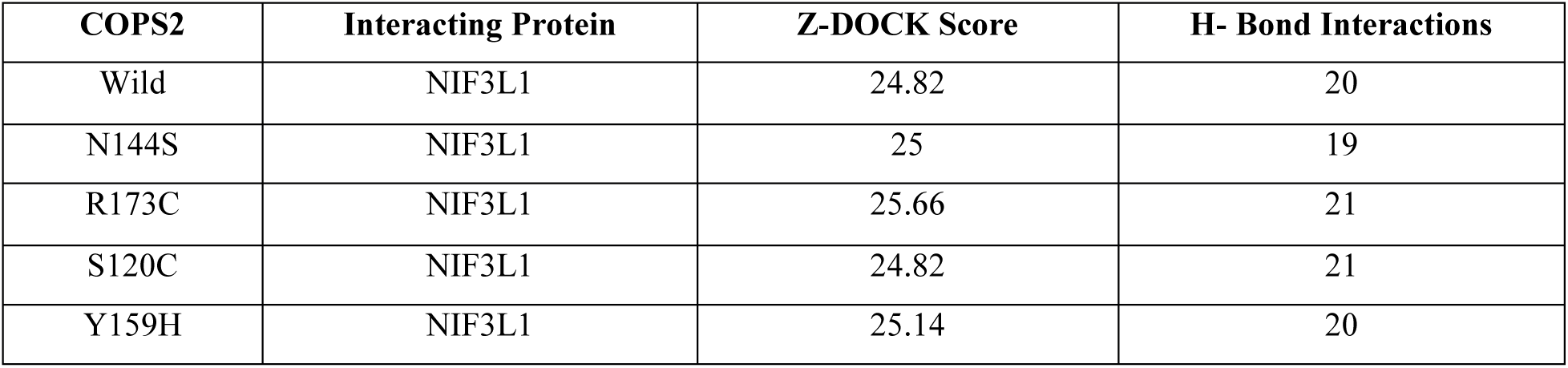
Protein-Protein Z-DOCK Score values of COPS2 Wild and mutant with NIF3L1 proteins models

These mutations resulted in best ZDOCK score which was further analyzed for monomer stability at a dynamic level. Further, the 3 nsSNPs (N144S, Y159H, R173C) [37–39] were submitted for Molecular dynamic simulation. The final screened nSNPs and wild structure are depicted in Figure 4

**Figure 4.**
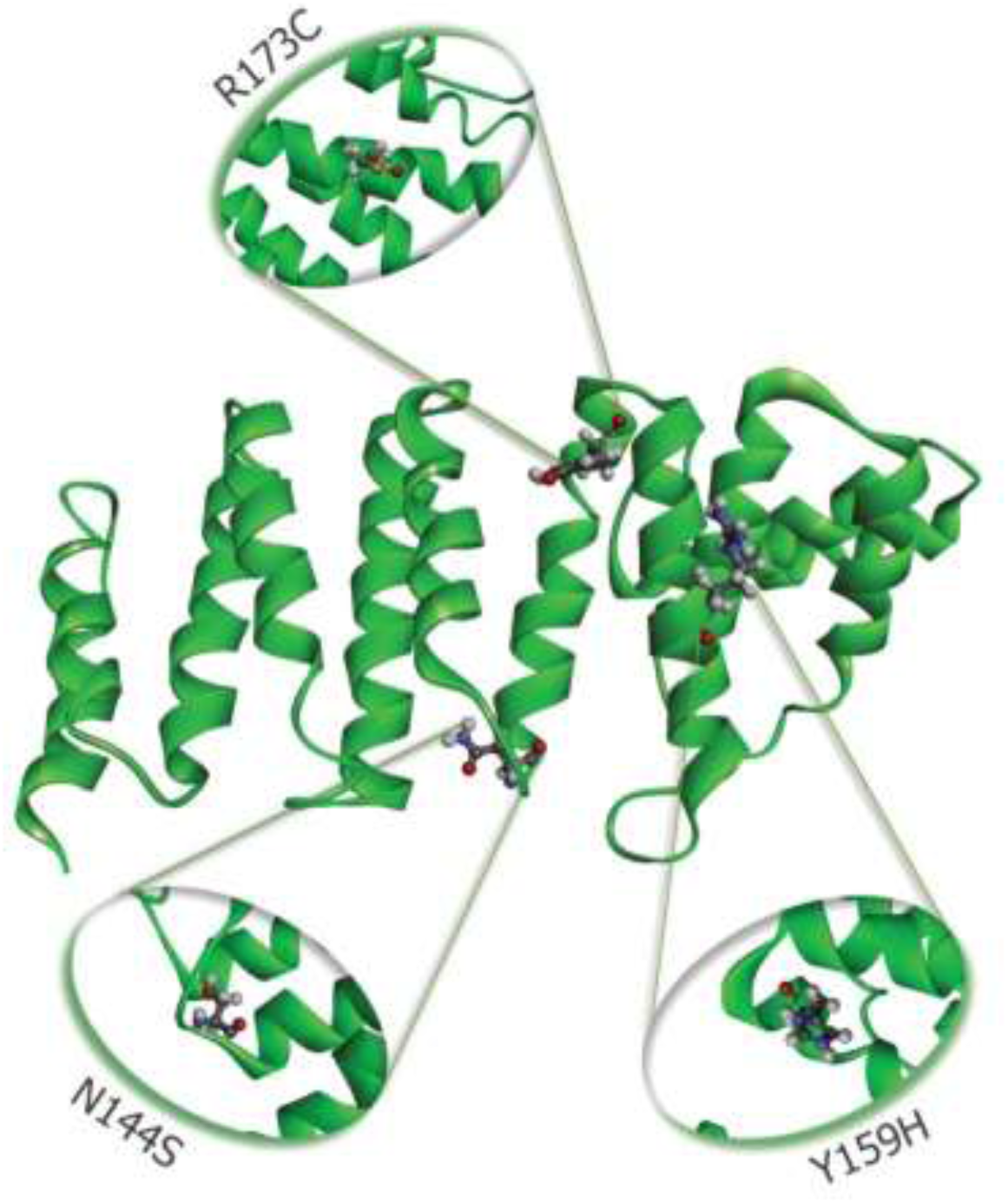
Schematic structural representation of missense variants (N144S, Y159H, R173C) location and details of alpha helices of COPS2 protein 3D structure.

### Molecular Dynamics Simulation

In order to explore the dynamic behavior of monomer (COPS2) with the three (N144S, Y159H, R173C) screened deleterious mutations in protein, molecular dynamic simulations were carried out for 150 ns at 5.4 pH. The Structural parameters analyzed were Root mean square deviation (RMSD), Root mean square fluctuation (RMSF), Radius of gyration (Rg), Number of hydrogen bonds (NH) and secondary structure (SSE) variation between WT and mutant structures N144S, Y159H and R173C. In order to bring back an all atoms level of detail, triplicate molecular dynamic simulation was carried for WT and mutant structures (N144S, Y159H, R173C). No such eloquent deviation in amino acid trajectories initiated in WT and in mutant structures while triplicating the MD simulation. The protein structures aligned with root mean square deviation (RMSD) values for Cα atoms were similar for all three trajectories and system for each dynamics of the individual structure showed similar RMSD value when compared with three trajectories. For initial 45ns there was considerable change in RMSD as protein requires equilibration so deviations were observed, for further analysis second half of the trajectories are considered after 45ns as the system becomes stable. After 45 ns the mutant structure (N144S) showed different deviation pattern Figure 6 (A) until the end of simulation resulting in Cα RMSD of 1.4 Å (Average:5.385,Standard Deviation: ± 0.970) drift from the WT(Average:4.698,Standard Deviation:±0.463) Whereas Y159H shows a vast deviation Figure 6 (B) with RMSD of 1.8 Å(Average:6.524,Standard Deviation:±0.971) drift from WT and R173C shows considerable deviation Figure 6 (C) of 0.3 Å(Average:4.797, Standard Deviation::± 0.760) drift from WT which explains structure instability as compared to WT RMSD as shown in Figure 6.

**Figure 5.**
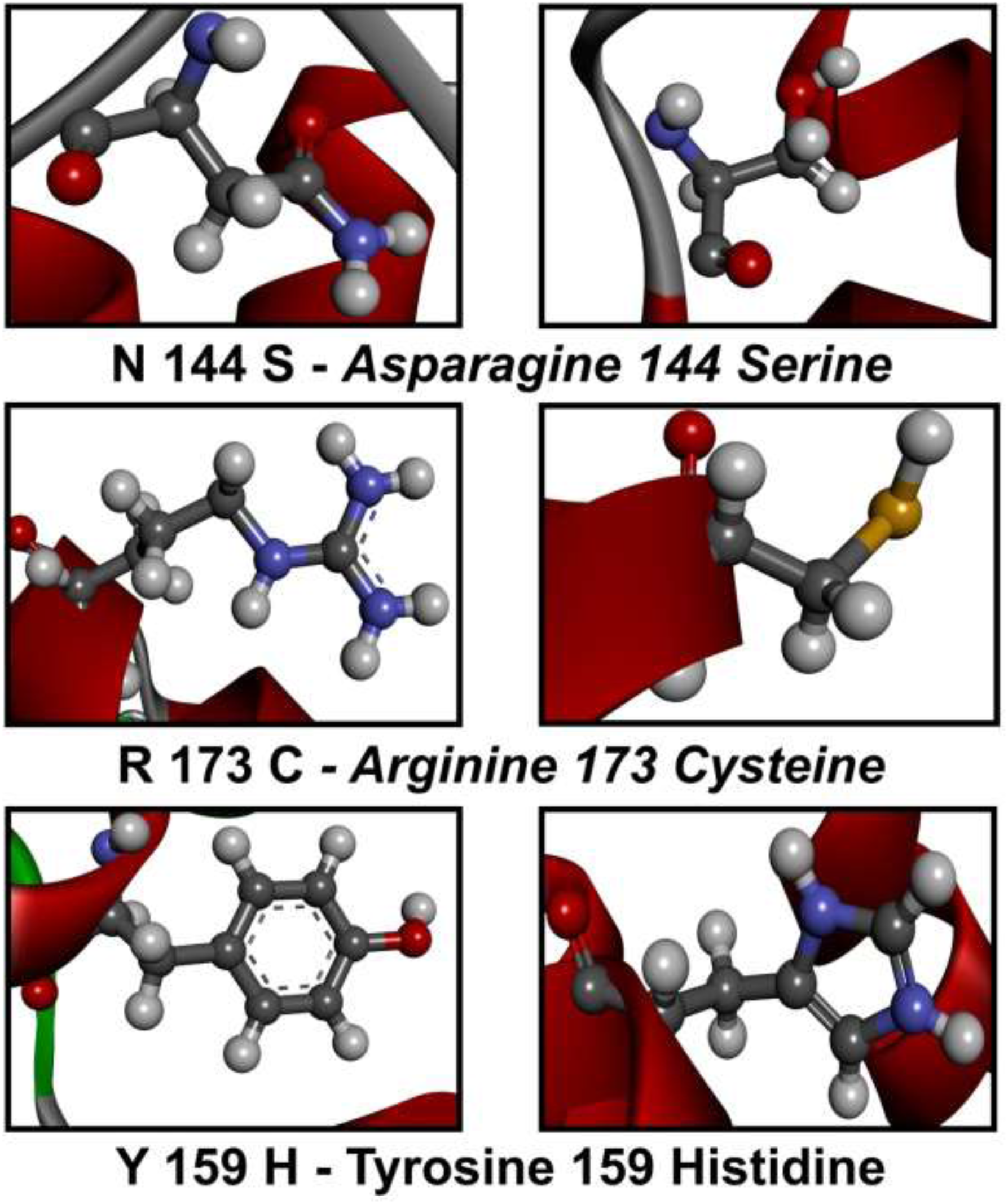
Schematic structural variation representation of mutations (N144S, Y159H, R173C) with corresponding comparison with wild type.

**Figure 6.**
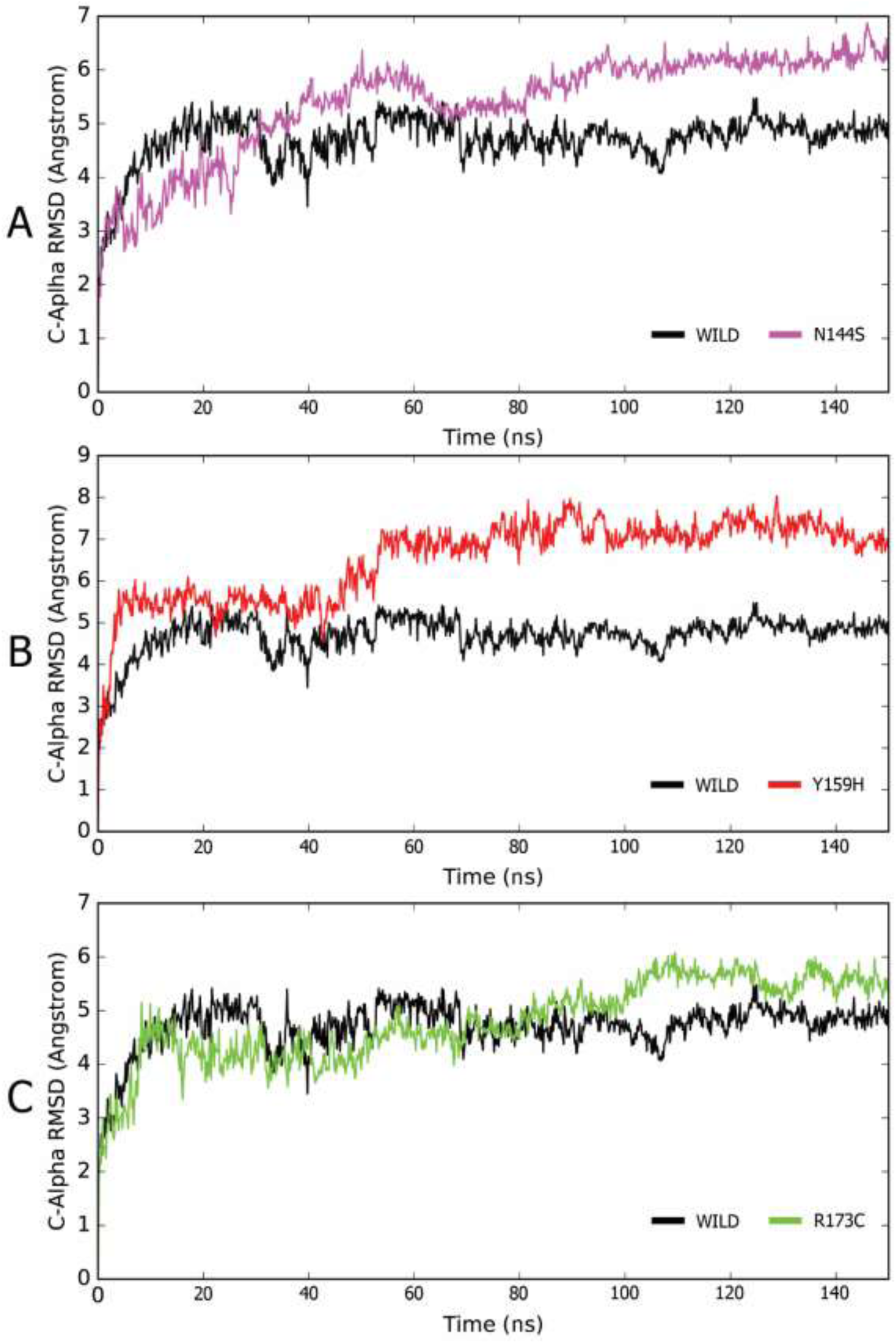
Root Mean Square Deviation graph for COPS2 mutant. A-(WT and N144S), B-(WT and Y159H) and C-(WT and R173C)

At 150 ns with instability as per RMSD graph shown with wild type, further investigation was carried to intervene in the flexible regions of protein structure and dynamics. Root Mean Square Fluctuation (RMSF) of Cα residues from its time-averaged position was calculated. RMSF explanation of residue Cα shows a higher fluctuation in mutant protein Y159H (Figure 7B) when compared to WT and the other mutant structures (N144S, R173C) during the period of the simulation. In Figure 7 (RMSF) with results of Y159H mutation affects nearby residues at a maximum of 5.3 Å, other two mutations N144S Figure 7 (A) and R173C Figure 7 (C) showing fluctuations at residue positions in Cα backbone 2.0 Å and 2.15 Å respectively as well as affecting the neighboring residues at maximum fluctuation of 3.4 Å and 4.2 Å respectively. When compared with WT in Figure 7. indicates an alteration in the flexibility of protein structure due to mutation.

**Figure 7.**
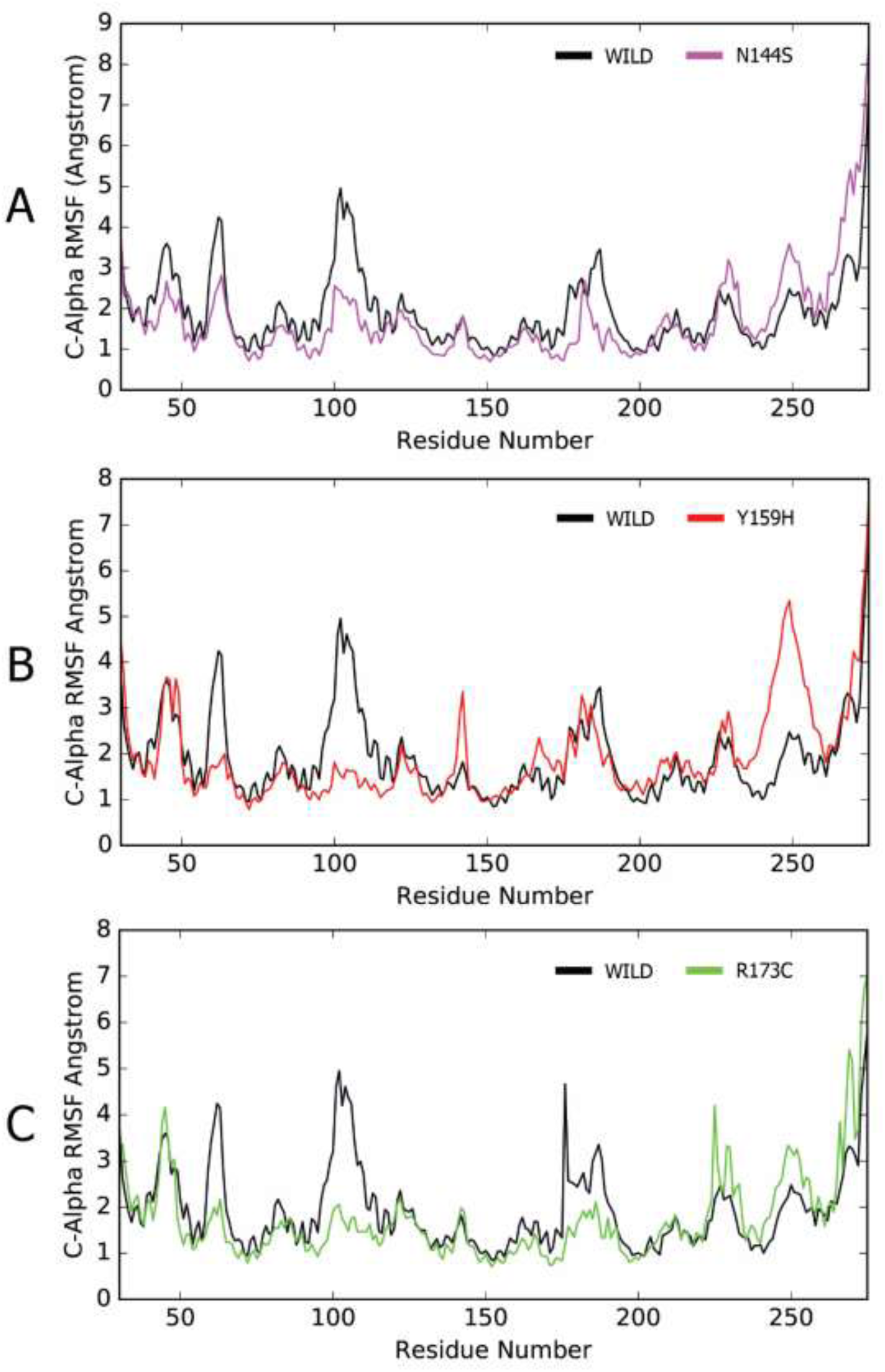
Root Mean Square Fluctuations (RMSF) Graph of COPS2 mutant in A-(WT and N144S), B-(WT and Y159H) and C-(WT and R173C)

In all these results predicts that there exists a eloquent change of structural deviation in mutant structures (Y159H, N144S, and R173C) when compared to WT. Additional information on structural flexibility is offered by secondary structural elements by the analysis of time-dependent secondary structure fluctuations of WT and mutations. The entire active domain consists of α-helix which shows stability in terms of WT structure with an α-helix content of 60.19% which concludes a total of 60.19% SSE content. Whereas in mutant model Y159H the α-helix percentage decreased to 59.07%, which indicates instability of protein structure for other two mutations N144S and R173C shows 60.09% and 60.97% respectively which has an impact on mutant model stability with each residue involved in altering the helix content. Secondary structure fluctuations are supported by forces such as H bond, which takes to further analysis of calculating the total number of H-bonds within the entire simulation, where the deviation in hydrogen bonds was recorded for WT and mutations, given in Figure 8. At Time series (150ns), where WT showed H-bond interactions with a range in (0 to 8) at 144 residue with respect to N144S it showed a range of (1 to 6) represented in Figure 8 (A). The alteration in hydrogen bonds, for WT at 159 residues the number of H-bonds deviations were in range (1 to 8) and for the mutation Y159H, it showed deviations ranging from (2 to 7) shown in Figure 8 (B). WT on 173 residue its shows deviations from range (0 to 5) so as for mutation R173C (0 to 6) shown in Figure 8 (C) which contributes to destabilization of the structure due to alterations in the number of H-bonds.

**Figure 8.**
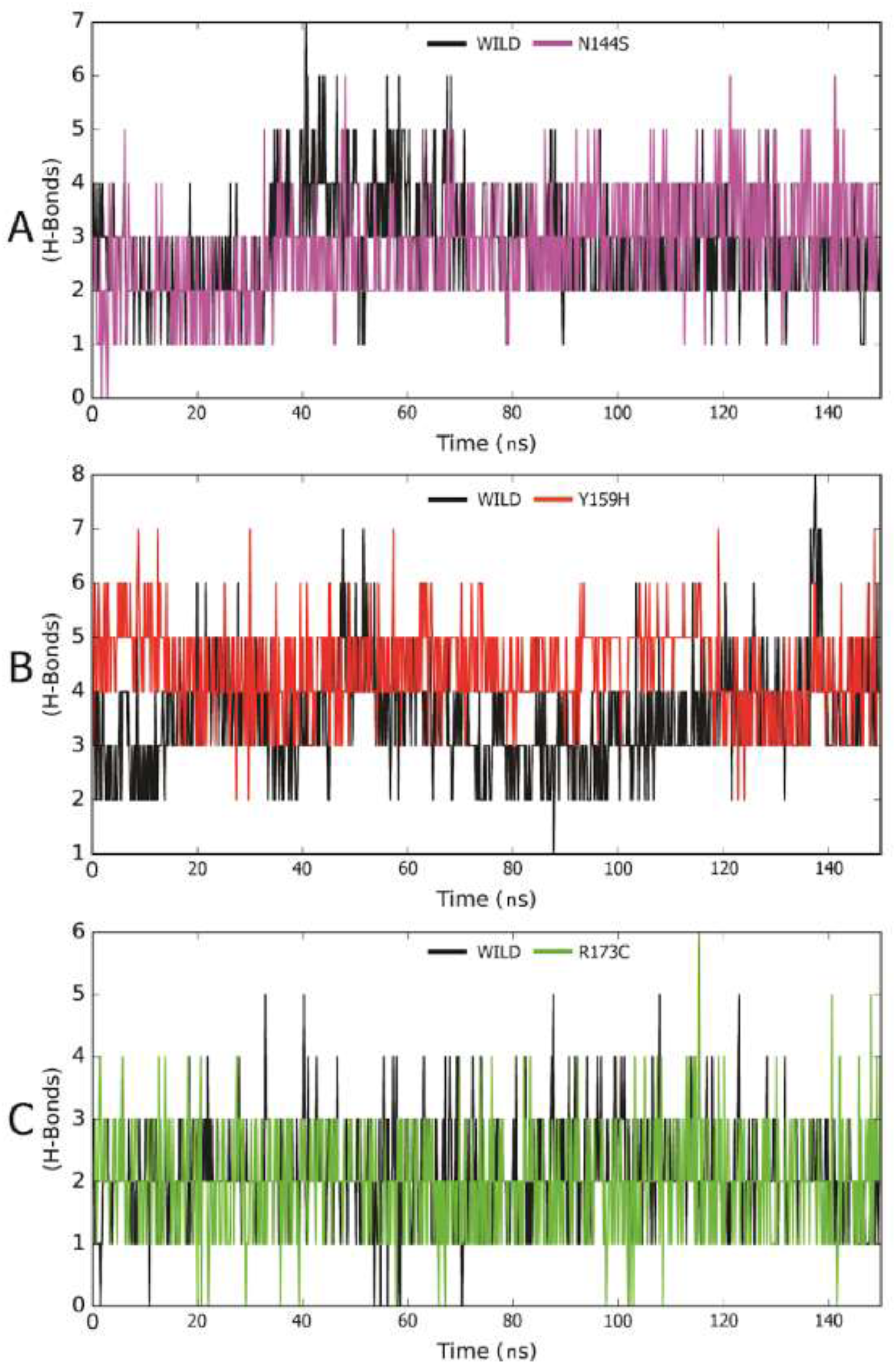
Number of Hydrogen bonds in with a comparison of WT with COPS2 mutant, A-(WT and N144S), B-(WT and Y159H)and C-(WT and R173C)

The atomic-level analysis which undergoes to investigate the radius of gyration explained as the mass-weighted root mean square distance of atoms from their center of mass. The entire simulation there was a deviation in the mutant models recorded as compared to WT, during the initial 45ns WT and mutation exhibits similar Rg value, after which mutant model (N144S) shows a deviation of ~0.8 Å from WT Figure 9 (A). Whereas Y159H shows the deviation of ~1.6 Å from the WT in Figure 9 (B) and when compared R173C with WT it shows a deviation of ~1.1 Å Figure 9 (C) which in respect to explains about the structure differences at an atomic level causing instability. The compactness of protein observed with mutant models states the deformity and which could harness the working efficiency of protein.

**Figure 9.**
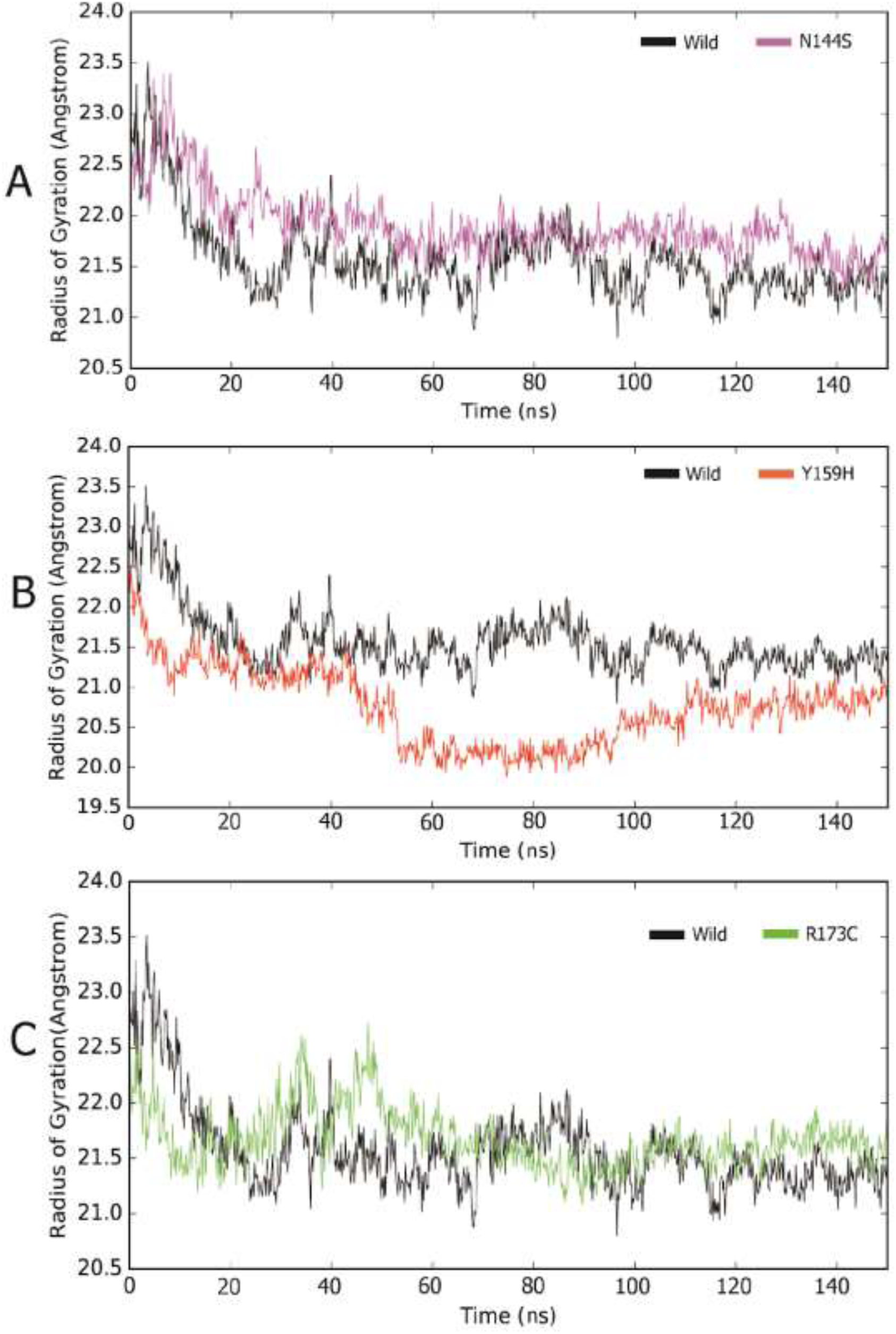
Radius of Gyration graph for COPS2 mutants in comparison with WT, where A-(WT and N144S), B-(WT and Y159H) and C-(WT and R173C) comparison at time series with Gyration (Å)

## Discussion

Amino acids substitution at specific position creates the structural level deviation affecting the chemistry of protein [Figure 5]. 90 non-synonymous coding SNPs were analyzed in search of deleterious nsSNP’s using function based prediction (SIFT, PolyPhen 2.0, PredictSNP, PROVEAN, and PMut)and stability based prediction tools (I-Mutant 2.0 and MuPro) for COPS2 gene. Genome identity in different individuals is around 99% and there is a change of 1% which includes 500 to 1000 base pairs inherited from generations and passed on concluding to diversity. The terminology for these changes or substitution is Single Nucleotide Polymorphism (SNP) which has a potential impact on diseases, drug competency, misregulation of protein expression, protein stability etc. Presence of nonsynonymous missense SNP’s in the coding region directly involved in the alteration or disrupt the normal functioning of the protein. As per the recent studies, gene *COPS2* is involved in neuronal differentiation and it is negatively regulated due to the transcription co-repressor domain interaction of NIF3L1 with COPS2. In which transcriptional corepressor is served by COPS2 component of COP9 signalosome (<u>Akiyama et al. 2003</u>). In order to intervene in structural changes of monomer stability for COPS2, a systematic approach towards mutational analysis was carried to observe the depth of nsSNP’s screened after function and stability based prediction tools resulting in 4 nsSNP’s (S120C, N144S, Y159H, R173C). Diving into the positional effects of mutations where Asparagine to Serine at 144 position affects the protein structure by removal of keto and amine group contributes to change in the chemistry of protein on the other hand where Tyrosine is substituted to Histidine where aromatic ring is lost and modulation of histidine with protein structure alters the stability of protein, taking in account where Arginine is replaced with cysteine alkyl side chain with amino triad is lost to sulphur which could give rise to disulphide bridges which collectively may change the geometric pattern causing deformity in protein structure.. All parameters for mutations resulted in a deviation from native structure which may conclude for causing structural instability. Our results demonstrate that the nsSNP’s we found are pathogenic and can lead towards an opportunity for the neural diseases association. During our analysis we also encountered some limitations with the computational study and computational facility and SNP database. For simulation of larger complexed protein molecule (COPS2-NIF3L1) for better understanding of COPS2 and NIF3L1 interactions in dynamic environment and effects of pathogenic mutations interaction and structural behavior with NIF3L1 in dynamic environment. The dbSNP database have serious limitations for studying complex disorders with an ethnic-dependent background due uneven and incomplete representation of the candidate SNPs in the databases for the major ethnic groups. Another limitation to in-silico analysis is completely based on different tools and their algorithms, which needs to be validated using in-vitro functional analysis.

## Conclusion

*Insilico* research has changed the fate of research with an innovative way of approaching biology with the help of computational algorithm in order to dive deeper into the disadvantages of nsSNP for structural alterations. Our study explained the pathogenicity of nsSNP’s by combining various sequence and structure principle sources predicting the deleterious SNP’s. Asparagine at 144 positions comes under conserved region reported in many studies and serves as lethal in substitution with Serine and also two other nsSNP’s Y159H and R173C describing the mutational effects in dynamic behavior with MD Simulation studies with RMSD, RMSF, SSE, NH Bonds and Rg analysis demonstrated insights of stability loss. As predicted destabilization of structure consequent could lead to functional failure of domain operation and can change the interaction behavior with the NIF3L1 protein. Change in the dynamic behavior of COPS2 nsSNP’s on active domain could lead to arrest the cell growth in G2 phase. Our study reported on these deleterious nsSNP’s could serve as pathogenicity and may create a opening room for neural diseases association.

## Conflict of Interests

The authors declare that there is no conflict of interests regarding the publication of this paper.

## Acknowledgments

The authors wish to thank the Management, Principal, Director, and Head of the Department of Biotechnology, Siddaganga Institute of Technology, Tumkur, Karnataka, India. The authors also appreciate KBITS for their support in providing them with the required computational resources for carrying out this project.

## Notes

### Competing Interest Statement

The authors have declared no competing interest.

